# Measuring antimicrobial use on dairy farms: a longitudinal method comparison study

**DOI:** 10.1101/2020.04.10.035485

**Authors:** G. M. Rees, D. C. Barrett, F. Sánchez-Vizcaíno, K. K. Reyher

## Abstract

Antimicrobial use on UK dairy farms is measured for surveillance purposes and utilizes veterinary sales data as a proxy for use. Two other methods of recording use have been used commonly on-farm: medicine waste bins and farm medicine records. However, none of these methods have been validated to measure antimicrobial use. The objectives of this research are to assess agreement between the 3 most common methods for measuring on-farm antimicrobial use with a pre-determined “gold standard” measure. Antimicrobial use was measured prospectively on 26 UK dairy farms using medicine waste bins into which participants placed all discarded medicine packaging for a 12-month period. At the end of 12 months, farm medicine records and veterinary sales data were obtained retrospectively for participating farms. The systematic difference between the mean on-farm antimicrobial use measured by each of the 3 methods with a gold standard measure was investigated using one-way repeated measures ANOVAs. Reliability and clinical relevance of the agreement between each pair of methods was quantified using the concordance correlation coefficient and the Bland-Altman method, respectively. Veterinary sales data shows excellent reliability for all forms of antimicrobial when compared with the gold standard. Medicine waste bins show moderate to excellent reliability for injectables, poor to good reliability for intramammaries and no agreement for other forms of antimicrobial. Farm medicine records do not show agreement for any form of antimicrobial when compared with the gold standard. The use of veterinary sales data as a proxy for on-farm antimicrobial use in the UK represents excellent statistical reliability and offers a clinically acceptable agreement with a gold standard method when used to measure both injectable antimicrobials and intramammary antimicrobials. These results have policy implications both nationally and internationally and are essential in quantifying the actual impact of agricultural antimicrobial use on both animal and human health.

## INTRODUCTION

Measuring antimicrobial use (**AMU**) is challenging (WHO, 2015; O’Neill, 2016; Kallen et al., 2019), and this is particularly true when measuring agricultural and veterinary AMU (RUMA, 2017; Mills et al. 2018; VMD, 2019a). An accurate understanding of AMU in animal health is essential for understanding patterns of resistance and informing antimicrobial stewardship policy from a One Health perspective (O’Neill, 2016). Indeed, the latest UK Government action plan specifically advocated “a clear need for more robust data on how antimicrobials are used to improve our understanding of the links between animal health and welfare, productivity, drug usage and resistance and to provide the evidence we need to design effective interventions and controls” (UK Government, 2019).

The UK measures veterinary AMU at a national level and publishes an annual report (VMD, 2019a), along with joint One Health reports with Public Health England (VMD, 2015; 2019b). The most recent One Health report shows veterinary AMU accounted for 36% of total UK use in 2017, although it has been acknowledged that AMU surveillance in agriculture is complex and current data are lacking validation (RUMA, 2017; VMD, 2019). Use of antimicrobials in the dairy sector has fallen by 30-35% since 2015, primarily through voluntary stewardship (RUMA, 2019; VMD, 2019a). In food-producing animals, dairy cattle represent the fourth highest user of antimicrobials by total weight (4.9 tonnes), behind pigs, poultry and gamebirds (VMD, 2019a). Since 2016, the data used to measure AMU in dairy cattle for the annual UK Veterinary Antimicrobial Resistance and Sales Surveillance report has estimated veterinary practice sales data as a proxy for use. These data are obtained from a small sample of UK veterinary practices and their representativeness is currently unknown. Pig and poultry sector AMU data, however, are considered robust, while AMU data from the beef and sheep sectors are currently in the process of being established (RUMA, 2015; 2019). Moving towards species-specific AMU data increases the granularity of such data, however the use of veterinary sales data as a proxy for use has not been validated (Mills et al., 2018; VMD, 2019).

The 3 most common methods for measuring on-farm AMU are veterinary sales data, on-farm medicine records and on-farm medicine waste bin audits. This paper presents a method agreement analysis of these common ways to measure AMU in dairy cattle. By assuming a gold standard measurement of actual AMU *a priori*, all 3 individual methods could be compared with the gold standard and an initial estimate of the appropriateness of each method made.

## MATERIALS AND METHODS

This study gained ethical approval from the University of Bristol Faculty of Health Sciences Research Ethics Committee; reference number 33021.

### Recruitment and Data Collection

Dairy farms (n=27) from South West England and South Wales were recruited to a wider study through purposive maximum-variation sampling using a combination of direct approach, nomination by local veterinary practices or self-nomination. Further details of herd characteristics can be found in Online Supplements Table 1, and details of sampling for this study can be found in Rees et al. (2018). Sample size estimation for reliability calculations was based on two observations per subject because all 3 methods of measurement were compared separately with the gold standard. An expected reliability value of 0.9 and an acceptable lower limit of reliability width of the 95% confidence interval of 0.7 were used, which gave a sample size requirement of 18 farms. This was then corrected to 23 farms based on an expected drop-out rate of 20% (Walter et al., 1998). This was deemed to be acceptable as 27 farms were enrolled onto the original project and complete data was collected for 26 farms.

An initial medicine inventory and structured questionnaire were undertaken on each participating farm; data on medicine name, quantity, number of individual items, storage location and expiry date of all antimicrobials present, farm demographics, herd health and management protocols were collected. Medicine waste bins were placed on each farm, and participants were requested to dispose of all used or discarded medicine packs (bottles, tubes, packaging etc.) in these bins. Bins were collected every quarter (90 days +/− 20); the final visit and second medicine inventory was conducted at day 365 (+/− 3 days). Farm medicine records were obtained at the final visit either in written or electronic form depending on the farmer’s usual record-keeping format and veterinary sales data was requested retrospectively from each farm’s veterinary practice for the duration of the study period. In the UK, all prescription veterinary medicine sales data must be recorded and stored by the veterinary practice. These records were computerized in all instances, although the software used, and the format provided varied between practices.

For each participating farm, an individual medicine workbook was created using Excel (Microsoft Office 365, USA) listing every medicine listed on the Veterinary Medicines Directorate’s Product Information Database (VMD, 2018).The contents of medicine waste bins were sorted, counted and data entered into the workbook. 10% of bins were double counted by a second researcher. Veterinary sales data and on-farm treatment record data were sorted, cleaned and entered into the workbook.

### Defining a “Gold Standard”

Developing a “gold standard” for AMU was necessary to devise a comparator for the 3 methods of recording, none of which had previously been validated. While the term ‘gold standard’ is often understood to mean the true value, in medical statistics the gold standard can be described as “the diagnostic test or benchmark that is the best available under reasonable conditions” (Versi, 1992). For this study, the gold standard was determined *a priori* and in discussion with experts in epidemiology, data handling and farm animal science, taking into account the potential for over- and underestimation of true AMU for each measure (Table 1). We determined that the most appropriate gold standard for AMU which minimized the potential for over- and under-estimation was based on a corrected value of veterinary sales data, adjusted by taking into account the full inventory of veterinary sales during the period between the beginning and the end of the study:

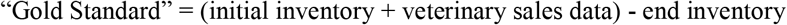

**Table 1.** Potential for over- or underestimation of antimicrobial use for the 3 different recording methods

| Method of measurement | Bias | Potential for error |
| --- | --- | --- |
| Veterinary sales data | Overestimation | - Purchased more antimicrobial during the time period than was used<br>- Wasted antimicrobials not accounted for<br>- Antimicrobials ascribed to wrong species |
|  | Underestimation | - Used antimicrobials which were purchased before the study period<br>- Antimicrobials purchased or obtained from sources other than veterinary practice<br>- Antimicrobials ascribed to wrong species |
| Farm medicine records | Overestimation | - Farmer records more antimicrobial being used than administered |
|  | Underestimation | - Farmer forgets or neglects to record treatments |
| Medicine waste bins | Overestimation | - Farmer discards medicines packaging into bins which were used before study period |
|  | Underestimation | - Farmer forgets or chooses not to use the bin or only uses the bin for some treatments and not others<br>- Antimicrobials used by the veterinary surgeon and not left on the farm |

The gold standard was based on veterinary sales data, as sales data were deemed to be least open to bias as they do not rely on farmer compliance or memory. The potential for a ‘time-lag’ in veterinary sales data could also be corrected for by taking a full inventory on the first and last days of the study. Therefore, while this gold standard was based in part on veterinary sales data, it is sufficiently different from that sales data to warrant comparison as it accounts for actual storage and use on farm.

Antimicrobials were classified according to their Veterinary Medicines Directorate classification for analysis as follows:

- Injectable antimicrobials: all antimicrobial products in injectable form
- Intramammary antimicrobials: all antimicrobial products in intramammary form
- Other antimicrobials: all antimicrobials that do not fit into the above two categories.

This included ocular preparations, tablets, boluses and powders used as footbaths.

Combination products containing at least 1 antimicrobial as an active ingredient were classified as antimicrobials. Methods of quantification of antimicrobials in the inventory and the medicine waste bins can be found in Rees et al. (2018).

### Data Analysis

Initially, one-way repeated measures ANOVAs were used to investigate whether there was a systematic difference between the mean on-farm antimicrobial use measured by 4 different recording methods for the following combinations: Veterinary sales vs. Gold standard, Medicine waste bins vs. Gold standard and Farm medicine records vs. Gold standard. Providing there is no evidence of a systematic difference between the measurements obtained from each pair of methods, the reliability and clinical relevance of the agreement between each pair was then quantified using the concordance correlation coefficient (**CCC**) and the Bland-Altman method, respectively. Analysis was conducted separately for the 3 different classifications of antimicrobials.

In the one-way repeated measures ANOVAs, “antimicrobial use” was the dependent variable and the independent variable was “recording method”. Where the normality assumption was not met after data transformation, a non-parametric Quade test was conducted to assess whether the distribution of values for each recording method was equal. The Greenhouse-Geisser correction was used where sphericity was violated. If significant results were found, post-hoc tests for differences between means were adjusted for multiple comparisons using Tukey’s test. When a Quade test was conducted, post-hoc tests for differences between distribution of values were adjusted for multiple comparisons following Holm’s method.

Reliability of methods was measured using CCC (Watson and Petrie, 2010). CCC point estimates along with 95% CIs were calculated using U-statistics (Carrasco et al., 2007). Values of CCC less than 0.5 indicate poor reliability, values between 0.5 and 0.75 indicate moderate reliability, values between 0.75 and 0.9 indicate good reliability, and values greater than 0.90 indicate excellent reliability.

The Bland-Altman method was used to get an insight into the pattern and extent of agreement between each pair of methods, as well as to determine whether such an agreement was likely to be clinically relevant at the farm level. This method calculates the ‘bias’ and 95% limits of agreement between 2 methods where the bias is the mean difference between the 2 methods (Bland and Altman, 1986).While the 95% limits show visually how well 2 methods of measurement agree, this quality judgement of this agreement depends on clinical context. The limits of maximum acceptable differences (limits of agreement expected) were defined prior to analysis, based on clinically and analytically relevant criteria agreed in discussions between the authors and clinicians working in dairy veterinary practice. Specifically, it was decided that if the 95% limits of agreement were within more than +/− 30% of the median total for the gold standard, this would equate to ‘clinically poor agreement’; within +/− 30% of the median total for the gold standard would represent ‘clinically reasonable agreement’; within +/− 20% of the median total for the gold standard would equate to ‘clinically good agreement’; and within +/− 10% of the median total for the gold standard would represent ‘clinically excellent agreement’. The influence of large outliers was evaluated by recalculating the limits of agreement with those outliers excluded (Watson and Petrie, 2010). Where the between-method differences did not follow a normal distribution, a logarithmic transformation of both measurements was conducted before analysis. If the normality assumption was not met after data transformation, a non-parametric form of the limits of agreement method was carried out as described by Bland and Altman (1999) and used instead for defining satisfactory agreement.

## RESULTS AND DISCUSSION

There is no evidence of systematic difference between the mean quantities of injectable antimicrobials (**INJAM**) and intramammary antimicrobials (**IMAM**) measured by the gold standard method and the mean amounts measured using veterinary sales data (INJAM: *P* = 0.995; IMAM: *P* = 0.999) or a medicine waste bin method (INJAM: *P* = 0.822; IMAM: *P* = 0.355), but there is a systematic difference with the mean amounts measured using the farm medicine records (INJAM: *P* < 0.001; IMAM: *P* = 0.04) (Online Supplementary Materials Table 2). In the case of other antimicrobials (**OtherAM**), mean quantities measured by the gold standard method are significantly different to the quantities measured by all the methods except for veterinary sales data (OtherAM: *P* = 0.47) (Online Supplementary Materials Table 2). Hence, veterinary sales are the only recording method for which both the reliability and the clinical level of agreement were evaluated for these OtherAM. Further information about the statistical tests and transformations used to investigate whether there was a systematic difference between the recording methods is shown in the Online Supplementary Materials Tables 2 & 3.

Based on CCC estimates, veterinary sales data show excellent reliability (95% CI >0.9) when measuring all 3 antimicrobial types (Table 2). In contrast, medicine waste bins show moderate to excellent reliability when measuring INJAM, and poor to good reliability when measuring intramammary antimicrobials (Table 2). Intraclass correlation coefficient (**ICC**) and CCC are 2 of the most popular overall indices used to assess agreement between methods when the outcome of interest is measured on a continuous scale (Carrasco and Jover, 2003). Both approaches are also advocated in Watson and Petrie’s (2010) review of the correct methodology for method agreement analysis. However, ICC is consistent only if the ANOVA model assumptions hold (Chen and Barnhart, 2008). In our study, the assumptions of normality and homogeneous variance were not met for some recording methods. Therefore, we used CCC instead, which was estimated using U-statistics, a recommended approach for skewed and non-normal data with low sample size (Carrasco et al., 2007).

**Table 2.** Concordance correlation coefficients (CCC) and statistical interpretation of reliability comparing veterinary sales and medicine waste bin data with a gold standard for different antimicrobial types

| Method comparison | Antimicrobial type | Concordance Correlation Coefficient (CCC) | 95% confidence intervals of CCC | Statistical Interpretation of CCC results |
| --- | --- | --- | --- | --- |
| Veterinary sales vs. “Gold standard” | Injectable antimicrobials | 0.998 | 0.996 – 0.999 | Excellent |
|  | Intramammary antimicrobials | 0.995 | 0.989 – 0.998 | Excellent |
|  | Other antimicrobials | 0.999 | 0.999 – 0.999 | Excellent |
| Medicine waste bin vs. “Gold standard” | Injectable antimicrobials | 0.821 | 0.620 – 0.921 | Moderate to excellent |
|  | Intramammary antimicrobials | 0.642 | 0.314 – 0.833 | Poor to good |

When measurements differed among methods, the Bland-Altman method was also used to determine whether those differences were likely to be clinically relevant at the farm level. Here veterinary sales data also show the best levels of agreement with the gold standard, with good to excellent agreement for INJAM and reasonable agreement for IMAM, although these data show clinically poor agreement for OtherAM (Table 3). In contrast, medicine waste bins show widely variable clinical agreement, ranging from poor to excellent agreement for INJAM and clinically poor to good agreement for IMAM (Table 3).

**Table 3.** Bland-Altman Plot statistics and clinical interpretation of agreement for all comparison of antimicrobial use recording methods

| Antimicrobial Type (unit) | Comparison | Mean difference (95% CI) | SD | 95% limits of agreement | Antilog of 95% limits of agreement | % farms within 95% limits | Assumptions met | Non-parametric 95% limits of agreement | Antilog of non-parametric 95% limits of agreement | % farms within non-parametric 95% limits | 95% limits of agreement compared with the median quantity recorded by the gold standard (based on non- | Clinical interpretation of agreement (based on non-parametric 95% limits) |
| --- | --- | --- | --- | --- | --- | --- | --- | --- | --- | --- | --- | --- |
| Injectable antimicrobials (ml) | Veterinary sales vs. gold standard | -118.11 (-255.03 to 18.80) | 338.89 | -782.34 to 546.11 | NA | 92.3 | Yes | NA | NA | NA | -14.21% to 9.92% | Good to excellent |
|  | Waste bin vs. gold standard | -999.63 (-2286.33 to 287.06) | 3184.89 | -7242.01 to 5242.75 | NA | 96.1 | No | -7938.75 to 425.62 | NA | 92.3 | (-144.14% to 7.73%) | (Poor to excellent) |
|  | Waste bin vs. gold standard<sup>1</sup> | -285.02 (-543.15 to -26.89) | 610.91 | -1482.4 to 912.36 | NA | 95.8 | No | -1618.5 to 450.38 | NA | 91.7 | (-30.8% to 8.57%) | (Poor to excellent) |
| Intramammary antimicrobials (tube) | Veterinary sales vs. gold standard | -0.519 (-30.94 to 29.90) | 75.30 | -148.10 to 147.07 | NA | 92.3 | Yes | NA | NA | NA | -21.53% to 21.38% | Reasonable |
|  | Waste bin vs. gold standard<sup>2</sup> | -0.13 (-0.31 to 0.05) | 0.43 | -0.98 to 0.72 | 0.37 to 2.06 | 96.2 | No | -0.99 to 0.12 | 0.37 to 1.13 | 92.3 | (-62.79% to 13.11%) | (Poor to good) |
|  | Waste bin vs. gold standard<sup>3</sup> | -0.05 (-0.09 to -0.01) | 0.096 | -0.23 to 0.14 | 0.79 to 1.15 | 96.0 | Yes | NA | NA | NA | -20.90% to 15.12% | Reasonable to good |
| Other antimicrobials (unit) | Veterinary sales vs. gold standard | -5.38 (-17.99 to 7.22) | 31.20 | -66.54 to 55.77 | NA | 92.3 | No | -69.25 to 51.06 | NA | 92.3 | (-153.89% to 113.47%) | (Poor) |

<sup>1</sup> two outliers removed
<sup>2</sup> log transformed
<sup>3</sup> log transformed and one outlier removed

For INJAM, veterinary sales data on average measure 118 ml less than the gold standard per farm over a 12-month period (Table 3 and Figure 1 (INJAM-a)). For 95% of farms, a yearly measurement of INJAM by veterinary sales data would be between 782.3 ml less and 546.1 ml greater than a measurement by the gold standard method (Table 3). Because these limits of agreement cross zero, veterinary sales data may under- or overestimate actual use of INJAM. This equates to a difference of 14.2% underestimation to 9.9% overestimation for 95% of farms when compared with the median total per farm measured by the gold standard method. The clinical interpretation of agreement is arguably the most important when comparing methods of measurement. Using the defined clinical agreement criteria, this represents good (within −20%) to excellent (within +10%) agreement between veterinary sales data and the gold standard method for INJAM (Table 3). Further results from the Bland-Altman method describing the agreement between veterinary sales and gold standard for IMAM are shown in Table 3 and in Figure 1 (IMAM-a), and for OtherAM in Table 3 and Figure 1 (OtherAM).

**Figure 1.**
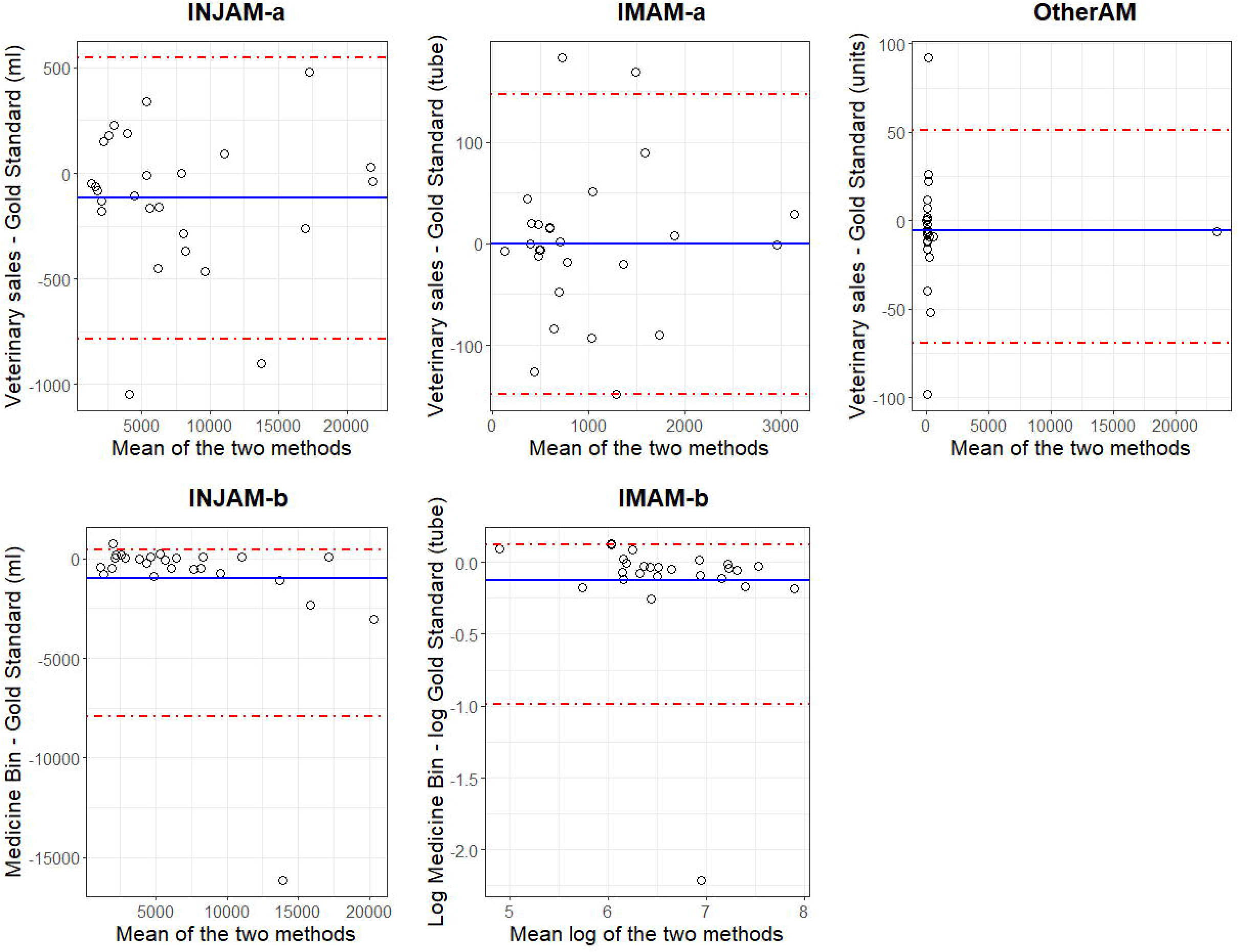
Bland-Altman plots comparing: INJAM-a: veterinary sales data with a gold standard recording method for injectable antimicrobials in total ml INJAM-b: medicine waste bin data with a gold standard recording method for injectable antimicrobials in total ml IMAM-a: Veterinary sales data with a gold standard recording method for intramammary antimicrobials in total number of tubes IMAM-b: Medicine waste bin data with a gold standard recording method for intramammary antimicrobials in total number of tubes OtherAM: Veterinary sales data with a gold standard recording method for ‘other’ antimicrobials in total number of units In each plot, the y-axis represents the difference between the two named methods and the x-axis represents the mean of the two methods; note that in IMAM-b, both y-axis and x-axis are in the logarithm scale. Dots represent the mean difference between the methods for each participating farm. The solid line represents the mean difference between methods across all farms. The hashed line represents the 95% limits of agreement in INJAM-a and IMAM-a; and it represents the non-parametric 95% limits of agreement in INJAM-b, IMAM-b and OtherAM. Abbreviations: INJAM – Injectable antimicrobials IMAM – Intramammary antimicrobials OtherAM – All other antimicrobials

Interestingly, the Bland-Altman plot reveals 2 large outliers where the gold standard method gives measurements for INJAM considerably above the medicine waste bin method (Figure 1 (INJAM-b)). Removing these outliers improves the closeness of the between-method differences to a normal distribution but does not solve the violation of normality. These 2 outliers also have a large influence on the mean difference between the 2 methods and on the limits of agreement, although not large enough as to change the clinical interpretation of the agreement (Table 3). A non-parametric form of the 95% limits of agreement show that the medicine waste bin method may produce values between 425.6 ml above the gold standard method to 7938.7 ml below the gold standard for INJAM (Table 3 and Figure 1 (INJAM-b)).

For IMAM, a logarithmic transformation of both the medicine waste bin measures and the gold standard measures slightly improves the closeness of the between-method differences to a normal distribution. A non-parametric form of the 95% limits of agreement derived from log-transformed data was back-transformed (antilog) to give limits for the ratio of measurements by these methods (Table 3) (Bland and Altman, 1986). The antilogs of these non-parametric 95% limits of agreement indicate that for 95% of farms the quantity of IMAM recorded by the medicine waste bin method would be between 0.37 and 1.13 times the quantity recorded by the gold standard method over a 12-month period. Thus, the medicine waste bin measurement may differ from the gold standard measurement by 63% below to 13% above actual use (Table 3). The Bland-Altman plot reveals a very large outlier where the gold standard method gives a measurement considerably above the medicine waste bin method (Figure 1 (IMAM-b)). Removing this outlier solves the violation of normality and has a substantial impact on the mean difference between the 2 methods and on the limits of agreement (Table 3). After excluding it, the medicine waste bin measurement differs from the gold standard measurement by 21% below to 15% above. Therefore, removing this outlier improves the clinical interpretation of the agreement between both methods for measuring IMAM from poor to reasonable. However, to the best of our knowledge the measurements captured by both methods for the outlier farm were correctly recorded and its removal cannot be fully justified. It is possible that this farm forgot to use the medicine waste bins, and this is important to capture as it may represent realistic use of this recording method. Some lack of agreement between different methods of measurement is inevitable. In this study, veterinary sales data tends to underestimate AMU but can both under- and overestimate use. Veterinary sales data differs substantially from the gold standard on certain farms; this can be explained by the fact that those farms either bought antimicrobials before the measurement period which were then used during this period or bought antimicrobials during this period which were not used until after measurement had ceased. However, these findings suggest that veterinary sales data is a valid method for measuring AMU which offers a clinically acceptable agreement with the gold standard method when used to measure both INJAM and IMAM. It is of note, however, that neither veterinary sales data nor the other alternative recording methods show clinically acceptable levels of agreement with the gold standard when measuring OtherAM. The reasons for this are not clear, but it may be that measuring OtherAM is complicated by the various units of measurements, depending on what pharmaceutical form the ‘other’ antimicrobial took. For example, ophthalmic ointments were measured on a per tube basis, while antimicrobial powders were measured per sachet, and tablets or boluses measured per packet. This presents difficulty when comparing these figures with those for INJAM or IMAM as the potential for over- or underestimation may vary by pharmaceutical form. These OtherAM are an important component of AMU surveillance; however, they make up a very small proportion of overall AMU, with INJAM and IMAM known to be the most commonly used and stored antimicrobials (Hyde et al., 2017; Rees et al., 2018; VMD, 2019a). Consequently, veterinary sales data may offer an acceptable alternative method for measuring AMU on dairy farms in the UK.

Medicine waste bins have been used in academic research to measure AMU on dairy farms in Canada and Peru (Saini et al., 2012; Redding et al., 2014; Nobrega et al., 2017). In this study, veterinary sales data outperformed medicine waste bin data as a proxy for actual on-farm AMU, due both to an increased reliability and a higher level of clinical agreement with the gold standard. Medicine waste bin audits were also more time-consuming, labor-intensive and required greater farmer acceptance and compliance. Thus, their use may be justified and potentially preferable in cases where obtaining veterinary sales data is difficult due to non-existence or data protection issues, but in most instances using veterinary sales data would be the superior method of estimating AMU.

In this study, farm medicine records were not a good method of measuring AMU as we have shown their mean measures to be statistically different to the gold standard for all antimicrobial types. It has previously been demonstrated that farmers place little value on maintaining accurate medicine records, see them as an unnecessary bureaucratic burden, deliberately omit certain medicines in order to achieve targets or forget to record medicines due to the practical constraints of medicine recording on a farm (Escobar, 2015). Improving the quality of farmer-recorded data (especially if access to good quality integrated electronic medicine records were available) could benefit AMU surveillance because such data benefits from increased granularity and chronology.

The use of veterinary sales data as a basis for calculating the ‘gold standard’ has obvious limitations. Comparing agreement between 2 methods where both rely on the same dataset makes it likely that the 2 methods will show some agreement. It is however clinically important not only to compare the 3 methods between themselves, but to attempt to validate these methods by comparing them with what is believed to be true (or the closest approximation to the truth). As such, each method needs to be compared with the closest approximate measurement to the true measurement, in order to be able to ascertain which method gives the best result. In this case, it was determined that the best possible gold standard was one based on veterinary sales data, for the reasons outlined in Materials and Methods. While this gold standard is based on veterinary sales data, the incorporation of on-farm data in the form of pre- and post-study inventories means the gold standard differs from veterinary sales data enough to justify comparing the 2. While it could be argued that the 2 methods not utilized when calculating the gold-standard are therefore less likely to compare well with the gold standard, this is still an important result. That veterinary sales data agrees best with the gold standard is not necessarily surprising, however validating its usefulness and importance can be of use to veterinary researchers and policymakers. That using medicine waste bins is a valid option for measuring injectable antimicrobials where veterinary sales data are not available or documenting that farm medicine records vary so greatly from the gold standard as to not be comparable are both important outcomes when considering methods to measure AMU.

The relatively small number of farms involved in this study mean that there is a risk of bias from outliers. Dairy farms contributing to this study were only located in the South West of the UK and were recruited purposively, so findings derived from this study cannot necessarily be considered to be generalizable. However, the characteristics of the recruited farms were broadly representative of the national picture (Online Supplementary Materials Table 3). Several assumptions were made when collecting and analyzing the data. Where labels on antimicrobials found in medicine waste bins had perished and the type of medicine was unidentifiable, these were disregarded. The proportion of medicine units this applied to was small (8/2809; 0.3%), meaning their exclusion from the study was unlikely to have affected overall conclusions. Where antimicrobials were in use or contained some remaining medicine, the quantity was estimated to the nearest 10%. Final inventory visits and bin collections were carried out on the 12-month anniversary of the study +/− 3 days. This led to a potential 6-day difference in the length of time some farms were studied, although it was assumed that this was unlikely to substantially affect the farm’s medicine recording given that for each farm the veterinary sales data and farm medicine records were measured for the same time period that bins were present.

## CONCLUSIONS

This study corroborates the use of veterinary sales data as a proxy for AMU on UK dairy farms. AMU data provided by medicine waste bins are inferior to that provided by veterinary sales data when compared with a gold standard and it is important to acknowledge and attempt to mitigate the current poor quality of farmer-recorded data identified in this study. Veterinary sales data is a valid method of recording AMU in the UK given that all veterinary antimicrobials are prescription-only, and that in general the veterinary surgeon both issues the prescriptions and supplies the antimicrobials. It should be noted that where prescription and supply is decoupled, for example where internet pharmacies are used or where legal decoupling of prescription and supply is proposed, veterinary sales data would not represent use.

## Acknowledgements

The authors would like to thank Prof. Henry Buller of the University of Exeter, Prof. Helen Lambert and Prof. Alastair Hay from the University of Bristol for their contributions to the wider project and also the participating farmers for their contribution to this research.

## Transparency Declaration

None to declare

## REFERENCES

Bland, J. M. and D. G. Altman. 1986. Statistical methods for assessing agreement between two methods of clinical assessment. Lancet 8(1):307–310. doi: https://doi.org/10.1016/S0140-6736(86)90837-8

Bland, J. M. and D. G. Altman. 1999. Measuring agreement in method comparison studies. Stat. Methods Med. Res. 8(2):135–160. doi: 10.1177/096228029900800204

Carrasco, J. L. and L. Jover. 2003. Estimating the Generalized Concordance Correlation Coefficient through Variance Components. Biometrics 59(4):849–858. doi: 10.1111/j.0006-341x.2003.00099.x

Carrasco, J. L., L. Jover, T. S. King, and V. M. Chinchilli. 2007. Comparison of Concordance Correlation Coefficient Estimating Approaches with Skewed Data. J. Biopharm. Stat. 17(4):673–684. doi: 10.1080/10543400701329463

Chen, C.-C. and H. X. Barnhart. 2008. Comparison of ICC and CCC for assessing agreement for data without and with replications. Comput. Stat. Data Anal. 53(2):554–564. doi: https://doi.org/10.1016/j.csda.2008.09.026

Escobar, M. 2015. DEFRA Report: Perceptions and practices of farm record-keeping and their implications for animal welfare and regulation. file:///C:/Users/gr14229/Downloads/12768_Finalreport-perceptionsandpracticesoffarmerrecord-keeping.pdf

Hyde, R. M., J. G. Remnant, A. J. Bradley, J. E. Breen, C. D. Hudson, P. L. Davies, T. Clarke, Y. Critchell, M. Hylands, E. Linton, E. Wood, and M. J. Green. 2017. Quantitative analysis of antimicrobial use on British dairy farms. Vet. Rec. 181(25):683. doi: 10.1136/vr.104614.

Kallen, M. C., S. Natsch, B. C. Opmeer, M. Hulscher, J. A. Schouten, J. M. Prins, and P. van der Linden. 2019. How to measure quantitative antibiotic use in order to support antimicrobial stewardship in acute care hospitals: a retrospective observational study. Eur. J. Clin. Microbiol. Infect. Dis. 38(2):347–355. doi: https://doi.org/10.1007/s10096-018-3434-0

Mills, H. L., A. Turner, L. Morgans, J. Massey, H. Schubert, G. Rees, D. Barrett, A. Dowsey, and K. K. Reyher. 2018. Evaluation of metrics for benchmarking antimicrobial use in the UK dairy industry. Vet. Rec. 182(13):379–379. doi: 10.1136/vr.104701.

Nobrega, D. B., J. De Buck, S. A. Naqvi, G. Liu, S. Naushad, V. Saini, and H. W. Barkema. 2017. Comparison of treatment records and inventory of empty drugcontainers to quantify antimicrobial usage in dairy herds. J. Dairy Sci. 100(12):9736–9745. doi: 10.3168/jds.2017-13116.

O’Neill, J. 2016. Tackling Drug Resistant Infections Globally: Final report and recommendations. The Review on Antimicrobial Resistance. https://amr-review.org/sites/default/files/160525_Final%20paper_with%20cover.pdf

Redding, L. E., F. Cubas-Delgado, M. D. Sammel, G. Smith, D. T. Galligan, M. Z. Levy, and S. Hennessy. 2014. Comparison of two methods for collecting antibiotic use data on small dairy farms. Prev. Vet. Med. 114(3-4):213–222. doi: 10.1016/j.prevetmed.2014.02.006.

Rees, G. M., D. C. Barrett, H. Buller, H. L. Mills, and K. K. Reyher. 2018. Storage of prescription veterinary medicines on UK dairy farms: a cross-sectional study. Vet. Rec. 184:153. doi: 10.1136/vr.105041.

RUMA Alliance. 2015. RUMA Guidelines: Responsible use of antimicrobials in cattle production. Responsible Use of Medicines in Agriculture Alliance. London, UK. https://www.ruma.org.uk/wpcontent/uploads/2015/07/RUMA_antimicrobial_long_cattle_revised_2015.pdf.

RUMA Alliance. 2017. Targets Task Force Report. Responsible Use of Medicines in Agriculture Alliance. London, UK. https://www.ruma.org.uk/wp-content/uploads/2017/10/RUMA-Targets-Task-Force-Report-2017-FINAL.pdf.

RUMA Alliance. 2019. Targets Task Force: Two Years On. Responsible Use of Medicines in Agriculture Alliance. London, UK. https://www.ruma.org.uk/wp-content/uploads/2019/10/RUMA-TTF-update-2019-two-years-on-FULL-REPORT.pdf

Saini, V., J. T. McClure, D. Leger, S. Dufour, A. G. Sheldon, D. T. Scholl, and H. W. Barkema. 2012. Antimicrobial use on Canadian dairy farms. J. Dairy Sci. 95(3):1209–1221. doi: 10.3168/jds.2011-4527.

UK Government. 2019. Tackling antimicrobial resistance 2019-2024: The UK’s 5-Year National Action Plan. Global and Public Health Group, ed. HM Government. London, UK. https://assets.publishing.service.gov.uk/government/uploads/system/uploads/attachment_data/file/784894/UK_AMR_5_year_national_action_plan.pdf

Versi, E. 1992. “Gold standard” is an appropriate term. BMJ (Clinical research ed.) 305(6846): 187. doi: 10.1136/bmj.305.6846.187-b

VMD. 2015. Joint report on human and animal antibiotic use, sales and resistance, 2013. UK One Health Report. Veterinary Medicines Directorate, ed, Addlestone, UK. https://assets.publishing.service.gov.uk/government/uploads/system/uploads/attachment_data/file/447319/One_Health_Report_July2015.pdf

VMD. 2018. Product Information Database. Veterinary medfìcines Directorate. in https://www.vmd.defra.gov.uk/ProductInformationDatabase/ Date accessed: 6th July 2017

VMD. 2019a. UK Veterinary Antibiotic Resistance and Sales Surveillance Report (UK-VARSS 2018). Veterinary Medicines Directorate. Addlestone, UK. https://assets.publishing.service.gov.uk/government/uploads/system/uploads/attachment_data/file/842678/PCDOCS-_1705145-v1-UK-VARSS_2018_Report_2019_FINAL_v2.pdf

Veterinary Medicines Directorate. 2019b. UK One Health Report: Joint report on antibiotic use and antibiotic resistance, 2013-2017. VMD, ed. Veterinary Medicines Directorate, Addlestone, UK. https://assets.publishing.service.gov.uk/government/uploads/system/uploads/attachment_data/file/775075/One_Health_Report_2019_v45.pdf

Walter, S. D., M. Eliasziw, and A. Donner. 1998. Sample size and optimal designs for reliability studies. Stat. Med. 17(1):101–110. doi: https://doi.org/10.1002/(SICI)1097-0258(19980115)17:1%3C101::AID-SIM727%3E3.0.CO;2-E

Watson, P. F. and A. Petrie. 2010. Method agreement analysis: a review of correct methodology. Theriogenology 73(9):1167–1179. doi: 10.1016/j.theriogenology.2010.01.003.

World Health Organisation. 2015. Global action plan on antimicrobial resistance. World Health Organization, http://www.who.int/iris/handle/10665/193736.

