## Supplementary materials for "Measuring antimicrobial use on dairy farms: a longitudinal method comparison study"

Online only data: Supplementary Materials

**Table 1:** Production, health and medicine storage characteristics of the 27 participating farms.

| **Property / characteristic** | **Median^1^ or Mean^2^*** | **Range (Median) or SD^4^ (Mean)*** |
| --- | --- | --- |
| Total herd size | **320** | *119 – 1,271* |
| Number cows in milk | **175** | *65 – 600* |
| Total annual milk volume (million L) | **1.1** | *0.5 – 4* |
| Total annual milk sales per cow (L) | **7500** | *3,600 – 11,300* |
| Milk price (pence per L) | **22.5*** | *15.3** |
| Somatic cell count (cells per ml) | **176,407*** | *50,466** |
| Bactoscan (1,000 bacteria per ml) | **20.9*** | *7.5** |
| Mastitis (cases per 100 cows per year) | **36.7*** | *15.9** |
| Lameness (cases per 100 cows per year) | **22.2*** | *15.3** |
| Respiratory disease (cases per 100 calves per year) | **10** | *0 – 47* |
| GI^5^ disease (cases per 100 calves per year) | **10** | *3 – 59* |
| Number PVM^3^ present on farm (by active ingredient) | **19** | *9 – 35* |
| Number PVM present on farm (by medicine unit) | **101** | *28 – 339* |

*^1^Median = non-normally distributed data*

*^2^Mean = normally distributed data^3^*

*^3^PVM = Prescription veterinary medicines*

*^4^SD = Standard deviation*

*^5^GI = Gastro-intestinal*

**Table 2**. Results of the ANOVAs, Quade test and post-hoc tests, assessing whether there was a difference between the mean on-farm antimicrobial use measured by three different recording methods when compared with a gold standard measure in a farm-level sample.

### **ANOVA results:**

INJAM: *F*_3,75_ = 12.91, *P* < 0.01, p-value after using Greenhouse-Geisser correction: *P*(GGe) < 0.01

IMAM: *F*_3,75_ = 3.268, *P* < 0.05, p-value after using Greenhouse-Geisser correction: *P*(GGe) < 0.05

### **Quade results:**

OtherAM: *F*_3,75_ = 16.224, P < 0.001

**Post-hoc results:**

| **Medicine type** | **Comparisons** | **Statistical method** | **Estimate**  **(95% Confidence Interval)** | **Standard Error** | ***P* value** |
| --- | --- | --- | --- | --- | --- |
| INJAM^1^ | Vet sales  – gold standard | ANOVA | -0.0086  (-0.146 to 0.129) | 0.053 | 0.995 |
| INJAM | Waste bin  – gold standard | ANOVA | -0.0464  (-0.184 to 0.091) | 0.053 | 0.822 |
| INJAM | Medicine records  – gold standard | ANOVA | -0.2875  (-0.425 to -0.150) | 0.053 | **<0.001***** |
| IMAM^2^ | Vet sales  – gold standard | ANOVA | -0.0053  (-0.207 to 0.196) | 0.078 | 0.999 |
| IMAM | Waste bin  – gold standard | ANOVA | -0.1287  (-0.330 to 0.073) | 0.078 | 0.355 |
| IMAM | Medicine records  – gold standard | Quade test | -0.2059  (-0.407 to -0.005) | 0.078 | **0.04*** |
| OtherAM^3^ | Vet sales  – gold standard | Quade test | N/A | N/A | 0.467 |
| OtherAM | Waste bin  – gold standard | Quade test | N/A | N/A | **<0.001***** |
| OtherAM | Medicine records  – gold standard | Quade test | N/A | N/A | **<0.001***** |

^1^INJAM – Injectable antimicrobials

^2^IMAM – Intramammary antimicrobials

^3^OtherAM – All other forms of antimicrobial

*P < 0.05

**P < 0.01

*** P < 0.001

**Table 3.** Statistical tests and transformations used to investigate whether there was a systematic difference between the mean on-farm antimicrobial use measured by three different recording methods when compared with a gold standard measure in a farm-level sample.

| Medicine type | Test | Transformation used for modelling |
| --- | --- | --- |
| Injectable antimicrobial | Repeated measures ANOVA | log10 (x + 100) |
| Intramammary antimicrobial | Repeated measures ANOVA | log10 (x + 10) |
| Other antimicrobial | Non-parametric Quade test | None |
